# A fast and accurate method for detection of IBD shared haplotypes in genome-wide SNP data

**DOI:** 10.1101/042879

**Authors:** Douglas W. Bjelland, Uday Lingala, Piyush S. Patel, Matt Jones, Matthew C. Keller

## Abstract

Identical by descent (IBD) segments are used to understand a number of fundamental issues in genetics. IBD segments are typically detected using long stretches of identical alleles between haplotypes in whole-genome SNP data. Phase or SNP call errors in genomic data can degrade accuracy of IBD detection and lead to false positive calls, false negative calls, and under‐ or overextension of true IBD segments. Furthermore, the number of comparisons increases quadratically with sample size, requiring high computational efficiency. We developed a new IBD segment detection program, FISHR (<u>F</u>ind <u>I</u>BD <u>S</u>hared <u>H</u>aplotypes <u>R</u>apidly), in an attempt to accurately detect IBD segments and to better estimate their endpoints using an algorithm that is fast enough to be deployed on the very large whole-genome SNP datasets. We compared the performance of FISHR to three leading IBD segment detection programs: GERMLINE, refinedIBD, and HaploScore. Using simulated and real genomic sequence data, we show that FISHR is slightly more accurate than all programs at detecting long (>3 cM) IBD segments but slightly less accurate than refinedIBD at detecting short (~1 cM) IBD segments. Moreover, FISHR outperforms all programs in determining the true endpoints of IBD segments, which is important for several reasons. FISHR takes two to four times longer than GERMLINE to run, whereas both GERMLINE and FISHR were orders of magnitude faster than refinedIBD and HaploScore. Overall, FISHR provides accurate IBD detection in unrelated individuals and is computationally efficient enough to be utilized on large SNP datasets > 20,000 individuals.

## Introduction

Two haplotypes (homologous chromosomal segments of DNA) can be defined as being identical by descent (IBD) if they descend from a common ancestor without either haplotype experiencing an intervening recombination (Powell et al. 2010). Using this definition, IBD haplotypes are identical at all measured and unmeasured genetic polymorphisms except at sites harboring (typically very rare) mutations that arose on either haplotype since the last common ancestor.

The probability of two individuals co-inheriting an IBD haplotype from a common ancestor at a given location is a function of the number of generations (*g*) since the common ancestor: *P*(*IBD* | *g*) = 2^1-2*g*^. Thus, siblings (*g*=1) have a 0.5 probability of sharing a segment IBD from one of their common ancestors (one parent) at a given genomic location, cousins (*g*=2) have a 0.125 probability, second cousins (*g*=3) a .03125 probability, and so forth. Although this probability drops off rapidly as a function of generations since the common ancestor, when haplotypes are shared IBD, they can be quite long, even for distantly related pairs of individuals. Under Haldane’s (1919) model of recombination, the length of IBD haplotypes shared between two individuals is exponentially distributed with mean 100/2*g* centiMorgans (cM). Thus, although a pair of individuals sharing a common ancestor 15 generations ago is highly unlikely to share any IBD haplotypes from that ancestor, when they do, the expected length of the segment is ~3.3 cM. Given that the probability of two random individuals sharing at least one common ancestor within 15 generations is ~1 in even large, randomly mating populations (Keller et al. 2011), a large number of IBD shared haplotypes around this length exist in any group of ‘unrelated’ individuals of the same population.

IBD shared haplotypes in samples with no known pedigree relatedness have been used for genotype imputation (Kong et al. 2008; Setty et al. 2011), IBD mapping (Vacic et al. 2014), heritability estimation (Browning and Browning 2013), phase inference (Kong et al. 2008), and inference of population structure (Palamara et al. 2012; Soi et al. 2011). Such IBD shared haplotypes are typically inferred from long stretches of identical alleles in phased, whole-genome single nucleotide polymorphism (SNP) arrays, but accurate and efficient IBD detection from such data is difficult for several reasons. First, phase and SNP-call errors can split long IBD segments into two or more shorter segments or lead to artificial truncation of IBD segments.

Such splitting and truncating of IBD segments can lead to failure to detect a segment altogether, due to the segment being shorter than a prespecified length threshold or due to the fact that shorter segments have lower posterior probabilities of being IBD, depending on the IBD detection algorithm. Thus, errors in SNP calling and phasing inflate false negative (miss) rates of IBD detection. Second, the sheer number of comparisons that must be made at each site (four comparisons between each pair of diploid individuals leads to a number of comparisons-twice the squared sample size), combined with the low base rate of true IBD segments between pairs of unrelated individuals, means that a substantial fraction of called IBD segments can be false positives. Similar to the case of false negatives, a false positive can be due to either an entire called segment not being IBD or to a called segment being overextended in one or both directions. Finally, because of the computational complexity of IBD detection, algorithms that sacrifice speed for accuracy can be unusable on the large sample sizes (e.g., >50,000) currently being accumulated (e.g., Schizophrenia Working Group of Psychiatric Genomics Consortium 2014; Sudlow et al. 2015). In a sample of 50,000 individuals, nearly 5 billion comparisons must be made per site. Thus, successful IBD detection programs must simultaneously meet a number of goals—computational efficiency, low false positive rates, low false negative rates, and accurate detection of IBD segment endpoints—that typically trade off with one another.

Several programs have been developed to discover IBD segments in SNP datasets when expected pedigree relatedness is low. GERMLINE (Gusev et al. 2009), often considered the benchmark IBD discovery program, is computationally efficient and therefore usable on very large samples, but the literature has indicated that its accuracy is lower than more recently developed programs. Because GERMLINE is fast and can be run in a way that leads to few false-negative calls at the expense of many false-positive calls, two newer IBD detection programs that reportedly outperform GERMLINE in accuracy, refined IBD (rIBD; Browning and Browning 2011) and HaploScore (Durand et al. 2014), use GERMLINE to detect candidate IBD segments. These candidate IBD segments are found using GERMLINE parameters that are optimized for each program. They are then post-processed, by extending, removing, or slicing the candidate segments in the hope of providing more accurate detection of IBD segments. rIBD uses a probabilistic hidden Markov model to give each candidate IBD segment obtained from GERMLINE a posterior LOD score as to whether it is truly IBD or not. rIBD has a lower false-positive rate than GERMLINE with only a modest increase in the false-negative rate, but it is computationally intensive and therefore has a very long runtime for large datasets. HaploScore uses information on the switch error rate and the SNP error rate to give a posterior probability of whether each candidate segment from GERMLINE is truly IBD or not.

The current paper describes a new program, FISHR (<u>F</u>ind <u>I</u>BD <u>S</u>hared <u>H</u>aplotypes <u>R</u>apidly), we developed to have a computational efficiency comparable to GERMLINE with accuracy as good as or better than rIBD or HaploScore. Importantly, because we had observed that existing programs tend either to over-extend true IBD segments or to split true IBD segments into multiple smaller ones, one of our central goals was to develop an algorithm that accurately determines the endpoints and hence the true lengths of IBD segments. This is important because bias in estimating the true length of IBD segments can lead to under‐ or over-estimates of heritability using IBD haplotypes, and inaccurate endpoint estimates can lead to decreased accuracy of imputation, phasing, and mapping near endpoints. As with rIBD and HaploScore, FISHR obtains candidate IBD segments by using GERMLINE. Segments can then be stitched together if separated by a small number of SNPs. After this, the number of “implied errors” (IE)—likely SNP call or phase errors—throughout the segment are counted, and the segment can then be shortened or removed entirely based on the number and location of the of IEs (see *Methods*). To analyze the programs, we compare the runtimes and offer extrapolated estimates for running them on large, whole-genome datasets. We then compare the positive predictive value (PPV, the proportion of called segments that are truly IBD) and sensitivity (the proportion of true IBD segments that are called) across a range of tuning parameters to explore the PPV-sensitivity trade-off for each program. We also compare the bias, precision, and accuracy of endpoint detection of truly IBD segments across programs and explain how these are related to PPV and sensitivity depending on how these metrics are defined. Much of the apparent discrepancy in comparisons of IBD detection programs that exist in the literature can be explained by how researchers have decided how over‐ and underextensions of called segments affect PPV and sensitivity.

## Results

### Comparison of run times

Figure 1 presents the log_2_ runtimes of the four programs as a function of sample size for five sample sizes. We calculated runtimes based on the optimal parameters found for each of the programs as described below. Runtimes were averaged from three separate simulated subchromosomes that were on average 16 cM long and contained 1,185 SNPs each (see *Methods*). Because GERMLINE is used as a first step for FISHR, HaploScore and rIBD, the runtimes for those programs include the time it took GERMLINE to find the candidate segments as well. Whereas GERMLINE is run internally for rIBD, FISHR and HaploScore require GERMLINE to be run separately and with user-specified parameters. Thus, in the present manuscript, we used three different sets of GERMLINE parameters: those that optimized accuracy for GERMLINE when reporting GERMLINE results, those that did so for FISHR for FISHR results, and those that did so for HaploScore for HaploScore results. For this reason, the runtimes presented in Supplemental Table 1 show different runtimes for GERMLINE when run by itself than when used as a precursor program.

**Figure 1.**
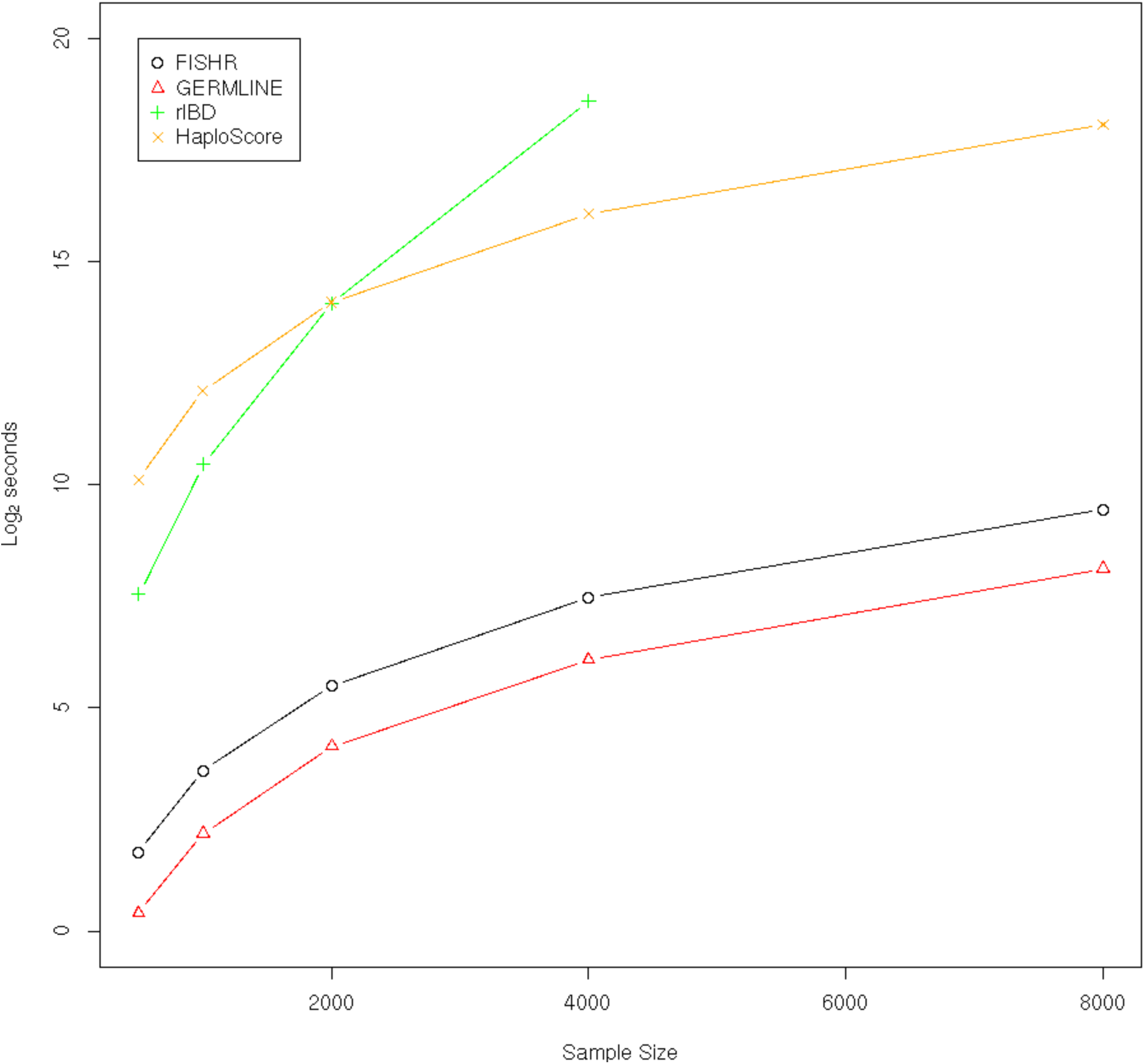
Runtime in log_2_ seconds for FISHR, GERMLINE, HaploScore, and rIBD at sample sizes of 500, 1,000, 2,000, 4,000, and 8,000 averaged from three 16-cM simulated chromosomal segments consisting of 1,185 SNPs each. rIBD with a sample size of 8,000 ran for one month (~2^21^ sec) before the server required maintenance and was shut down.

GERMLINE was the fastest program to run at any of the sample sizes, with FISHR doubling to quadrupling its runtime at all sample sizes. Most of the increase in runtime for FISHR compared to GERMLINE was caused by using a smaller minimum cM threshold for the initial GERMLINE segment discovery, which is necessary in order for FISHR to stitch together any segments that GERMLINE splits apart. Both HaploScore and rIBD had runtimes hundreds to thousands of times longer than FISHR, with this ratio increasing with larger sample sizes for rIBD. To gauge how the programs performed on a realistic, large SNP dataset, we also calculated runtime on a sample of 17,093 individuals aggregated from four datasets (the Atherosclerosis Risk in Communities cohort, the Coronary Artery Risk Development in Young Adults study, the controls from the Molecular Genetics of Schizophrenia study, and the GENEVA Genes and Environment Initiative in Type 2 Diabetes study; dbGap accessions phs000280.v2.p1, phs000285.v3.p2, phs000167.v1.p1, and phs000091.v2.p1, respectively) from the NIH Genotype and Phenotype database. Because IBD detection is typically done in parallel for each subchromosome arm, we analyzed the longest chromosome arm, 5q, which contained 19,772 SNPs on the Affy 6.0 SNP array. When the threshold for segment length was set to 1 cM, GERMLINE took about 1.5 days to run, FISHR took about 6.5 days (including 5 days, 16 hours for GERMLINE initial candidate segment discovery), whereas both rIBD and HaploScore ran for nearly two months before the server required maintenance and the processes were stopped. From extrapolations of the runtimes on simulated data (Figure 1), we predict that HaploScore would have finished running in just over two months and rIBD would have required over a year to finish.

### PPV and sensitivity in simulated data

PPV and sensitivity are the most common metrics in this literature for comparing the accuracies of the programs, and so we focus on these for commensurability. An inherent tradeoff exists between the two metrics: conservative calling algorithms that call fewer IBD segments tend to have relatively high PPVs and low sensitivities, whereas more liberal calling algorithms that call more IBD segments tend to have relatively high sensitivities and low PPVs. Figure 2 illustrates how we defined PPV and sensitivity depending on the degree to which called segments over‐ or underextend the endpoints of true IBD segments. For PPV, we first calculated the total length of overlap between each called segment and any corresponding true IBD segment(s) and divided this overlap by the length of each called segment. Thus, this proportion was 1 for the called segment in Figure 2A and for both of the called segments in Figure 2D, < 1 for the called segments in Figures 2B, 2C, and 2E, and 0 for called segments that did not overlap any true IBD segments. When a single called segment overlapped multiple true IBD segments (Figure 2E), overlap was defined as the sum of the overlapping lengths. PPV was then calculated as the average of these proportions across all called segments weighted by their length in basepairs. Similarly, for sensitivity, we calculated the length of total overlap between each true IBD segment and any corresponding called segment(s) and divided this overlap by the length of the true IBD segment. When multiple called segments split up a single true IBD segment (Figure 2D), overlap was again calculated as the sum of the overlapping lengths. Thus, these proportions were <1 for Figures 2A, 2C, and 2D but 1 for Figure 2B and for both true segments in Figure 2E. We defined sensitivity as the average of these proportions weighted by base pair length across all true IBD segments. Alternative definitions of these metrics are possible. For example, proportions greater than a threshold (.5) have been treated as true positives and those less than .5 as false positives for calculating PPV (Browning and Browning 2011). We prefer our definitions because they result in PPV and sensitivity being continuous functions, rather than step functions, of the degree of over‐ or underextension, respectively.

**Figure 2.**
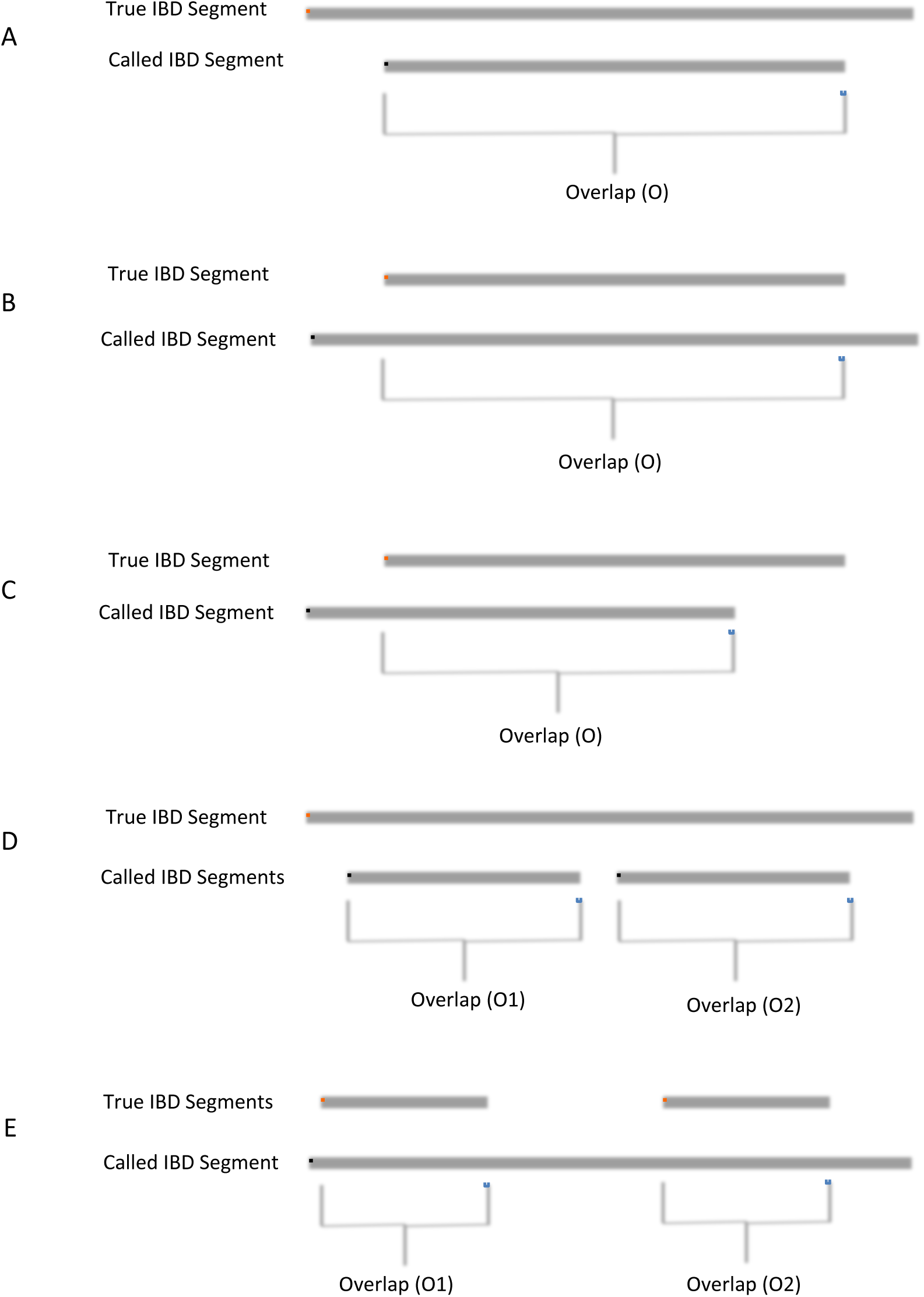
Method for calculating PPV and sensitivity from the called IBD segments and the known true IBD segment from an (A) underextended call, (B) overextend call, (C) off-center call, (D) situation where two called segments occur within a single true IBD, and (E) situation where one called segments occurs within two true IBD segments. For each called segment, we divided the length of the overlap with the true segment (O) or sum of the overlaps (O1+O2) by the length of the called segment. PPV was the average of these proportions across all called segments, weighted by base pair length. To determine sensitivity, for each true segment, we divided the length of overlap (O) or sum of the overlaps (O1+O2) by the length of true IBD segment. Sensitivity was the average of these proportions across all true IBD segments, weighted by base pair length. When two called segments overlapped one true IBD segment (D), two proportions contributed to PPV (one for each of the called segments) but one proportion to sensitivity. Conversely, when one called segment overlapped two true IBD segments (E), one proportion contributed to PPV and two to sensitivity.

To estimate the accuracies of the programs, we used perfectly matching phased haplotypes from simulated, dense sequence data with no phase or call errors to define the endpoints of true IBD segments (see *Methods*). We then called segments by applying each of the programs to a subset of the sequenced variants designed to mimic phased SNP array data, with realistic linkage disequilibrium (LD) patterns, allele frequencies, SNP densities, and levels of SNP-call and phase errors. Figure 3 displays PPV and sensitivity where both called and true IBD segments had minimum lengths of 3 cM (Figure 3A) or 1 cM (Figure 3B). For each program, we varied thresholds to produce a spectrum of conservative to liberal segment calling. In particular, we varied the *moving average* threshold for FISHR, the minimum *LOD score* for rIBD, and the *bits* argument for GERMLINE and HaploScore. At 3 cM minimum segment lengths, FISHR outperformed every other program with a higher PPV for any given sensitivity or, alternatively, a higher sensitivity for any given PPV. At 1cM minimum threshold lengths, FISHR and rIBD performed similarly and outperformed both GERMLINE and HaploScore.

**Figure 3.**
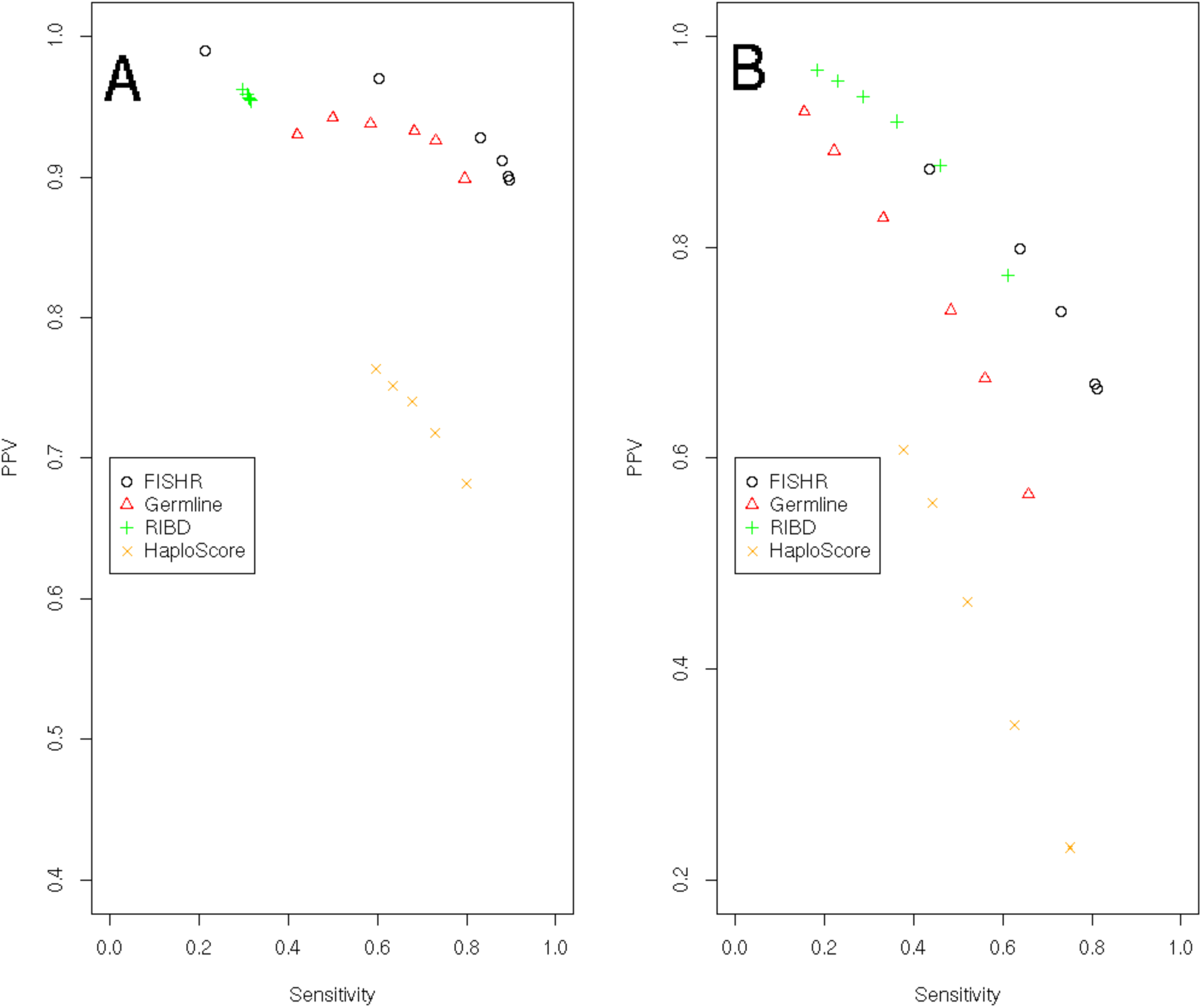
PPV-Sensitivity plots for FISHR (o), GERMLINE (Δ), rIBD (+), and HaploScore (x) when calculated using a minimum of 3 cM for called IBD and a minimum of 3 cM for true IBD (A) and when using a minimum of 1 cM for called IBD and a minimum of 1 cM for true IBD (B).

By using the same minimum-length thresholds (e.g., 3 cM) for both the called and true IBD segments, the results displayed in Figure 3 are highly sensitive to the accuracy of the endpoints of the called segments, as well as to truncation and splitting errors. For example, all sensitivity estimates of rIBD in Figure 3A are less than 0.3, below those of other programs and below those reported in the manuscript introducing rIBD (Browning and Browning 2011). As we demonstrate below, this is because rIBD tends to split true IBD segments into multiple, smaller called segments; when these called segments are shorter than the threshold (e.g., 3 cM), they are dropped for the purposes of calculating sensitivity, and therefore most true IBD segments > 3 cM appear to be missed. Because the endpoints of segments called by GERMLINE and especially FISHR are more accurate (see below), the performances of these programs are not degraded to the same extent. An alternative definition of PPV that is less affected by such truncation/split errors is to compare all called segments greater than a length threshold (3 or 1 cM) to all true IBD segments that are at least half that length (1.5 or 0.5 cM, respectively). Similarly, sensitivity can be computed by comparing all true IBD segments greater than 3 or 1 cM to all called segments greater than 1.5 or 0.5 cM, respectively. Figure 4 shows PPV and sensitivity calculated in this way. The performance of all programs improved but the improvement was greater for programs that are inaccurate at endpoint estimation (rIBD and HaploScore) than for programs that are more accurate at endpoint estimation (GERMLINE and especially FISHR; see results on endpoint accuracy below). At 3 cM minimum called (PPV) and true IBD (sensitivity) segment lengths, FISHR performed slightly better than GERMLINE or rIBD, whereas at 1 cM minimum thresholds, rIBD outperformed FISHR. Because rIBD uses a posterior probability instead of a minimum cM length threshold to call segments, Figure 4 also shows rIBD results when no minimum length was used in calculating sensitivity and when much smaller true IBD lengths (0.5 cM for Figure 4A and 0.25 cM for Figure 4B) were used for calculating PPV. The sensitivity values for these instances of rIBD were improved and show rIBD to be superior to all other programs with respect to IBD detection accuracy. However, as demonstrated above, these conclusions rest upon arbitrary decisions on how PPV and sensitivity are defined. Moreover, as demonstrated below, the improved sensitivity of rIBD when there was no minimum length of called segments occurred because rIBD often splits long, true IBD segments into multiple, short called segments, which were sometimes dropped when a length threshold was used in calculating sensitivity.

**Figure 4.**
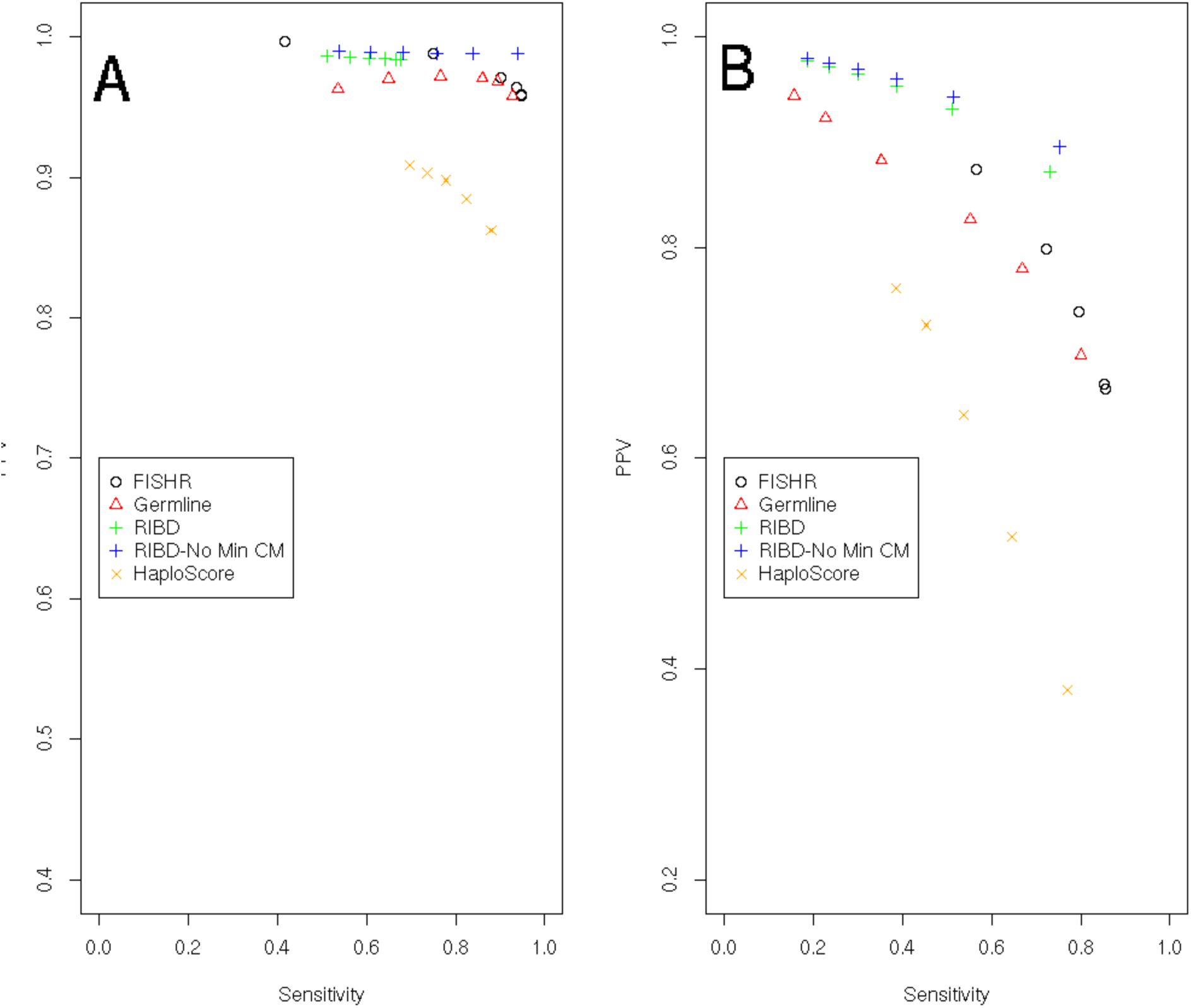
PPV-Sensitivity plots for FISHR (o), GERMLINE (Δ), rIBD (+), and HaploScore (x) when calculated using a minimum of 3 cM for called IBD and a minimum of 1.5 cM for true IBD (A) and when using a minimum of 1 cM for called IBD and a minimum of 0.5 cM for true IBD (B). Additional measures are present for rIBD (+) using a minimum true IBD length of 0.5 cM for PPV and no minimum called cM length for sensitivity (A) and a minimum true IBD length of 0.25 cM for PPV and no minimum called cM length for sensitivity (B).

### Accuracy of called segment endpoints in simulated data

As noted above, the differences between the results in Figures 3 and 4 correspond to how accurately the endpoints were estimated by each program. To quantify accuracy of endpoint estimation, we first found optimal parameters for each program by searching through combinations of the various input parameters, choosing those that maximized the sum of PPV and sensitivity. Using these parameters, we divided the length of over‐ or underextension of each called segment endpoint by the length of the corresponding true IBD segment. Figure 5 shows the distribution of these proportions—the degree to which each endpoint was over‐ or underextended—when called segments had minimum length of 3 cM and true IBD segments had minimum length of 1.5 cM (results for 1 cM called and .5 cM true thresholds are shown in Supplemental Figure S1). It should be noted that using a 3 cM threshold for called and 1.5 cM for true IBD segments was the optimal scenario for all programs (Supplemental Figure S2). Any called segment that had no corresponding true IBD segment (false positive) was given an arbitrary value of 1 and any truly IBD segment with no corresponding called segment (false negative) was given a value of ‐1. The text to the left of each histogram shows the bias (defined as the mean proportion), precision (defined as the standard deviation of the proportion), and accuracy (defined as the standard deviation from 0 rather than from the mean proportion) when the false positive and false negative calls were included. Accuracy provides an estimate of how accurate the called segments are compared to perfect calls with no under‐ or overextension, and incorporates information on both bias and precision (accuracy^2^ = bias^2^ + precision^2^). FISHR had the most accurate (0.227) endpoints and was the most precise (0.227) of all algorithms. FISHR also showed very little bias (-0.011) with respect to under‐ or overextending calls. HaploScore (bias = 0.077) tended to overextend segments, whereas GERMLINE (bias = ‐0.044) and to a greater extent rIBD (bias = ‐0.177) tended to call segments that were shorter than the true IBD segments. rIBD also tends to miss truly IBD segments at a much higher rate than either FISHR or GERMLINE while HaploScore tends to both miss true IBD segments and call segments which are not IBD, as shown by the large values at ‐1 and 1, respectively. These conclusions remained unchanged when we excluded false positive and false negative calls (reported on the right side of histograms in Figure 5).

### Accuracy of called segment endpoints in real data

All previous results used simulated data where the true IBD segment endpoints were known within a small margin of error. To determine how well the programs detect IBD segment endpoints in real data, we obtained data from 1,872 unrelated individuals from the UK10K dataset (The UK10K Consortium 2015), who were whole-genome sequenced at over 28 million markers. We extracted markers in the Illumina 650K SNP panel, re-phased them using SHAPEIT2 (Delaneau et al. 2012), and called segments from each of the four programs on this SNP dataset (see *Methods*). All remaining markers were retained as a holdout sample to calculate opposite homozygosity (OH) in and around regions where segments were called by each program. OH (e.g., an A-A genotype in one individual and a C-C genotype in the other) at masked markers within and around the called segments can be used to estimate the programs’ rates of false-positive and false-negative calls and to infer where called segments over‐ or underextended true IBD segments (Browning and Browning 2012). Even when the underlying haplotypes are truly IBD, sporadic mismatching alleles within a called segment can occur due to SNP errors, and a string of such mismatches can occur due to one or more phase errors.

However, phase errors cannot cause OH at true IBD locations; only the rare event of SNP call errors changing a heterozygous SNP to the opposite homozygous call can cause (very low levels of) sporadic false OH in the data. Therefore, locations where the rate of OH in holdout markers is high *within* the boundaries of called segments suggest regions of false positive calls (typically overextended segments), whereas locations where the rate of OH is low *outside* the boundaries of called segments suggest regions of false negative calls (typically underextended calls).

Figure 6 shows an example of a region where all four programs called a segment between two individuals and the locations where OH occurred in the holdout sequence data. To compare these instances of OH to the rate of OH expected in a pair of non-IBD segments, we also show the locations of OH at all holdout markers between a pair of randomly selected individuals at this location. Given the highly discrepant rate of OH between the focal pair and the rate of OH between the random pair, it is safe to assume that a true IBD segment existed between the focal individuals at this region, and the endpoints of this true IBD segment can be roughly inferred from where the OH rates between the focal individuals increase in the holdout sequence data.

The results depict a fairly typical example in which rIBD apparently broke up a long true IBD segment into multiple short called segments. FISHR, GERMLINE, and HaploScore appear to have done better in this example at discovering one long true IBD segment, with the main differences between programs being where the endpoints were estimated. Supplemental Figures S3-S22 display an additional 20 similar examples chosen at random from among 5 FISHR called segments, 5 rIBD called segments, 5 HaploScore called segments, and 5 GERMLINE called segments.

To quantify the accuracy of the called segment endpoints for each program in this real dataset, we calculated the proportion of OH (POH) of holdout markers in 4 quarters of each called segment from the UK10K data, as well as two regions of the same base-pair length upstream and downstream from the called segment. We then calculated the average POH of the four quaters and two quarter-length flanking regions for each called segment. These results are presented in Figure 7 and corroborate our earlier conclusions about endpoint accuracy of the four programs in the simulated data (Figure 5). Figure 7A displays the four quarters of the called segment and the flanking regions, whereas the Figure 7B displays only the first through fourth quarters within the called segments on an expanded scale. Figure 8 illustrates how these POH profiles should appear for programs that estimate endpoints perfectly, tend to underextend them, tend to overextend, or both. Of the four programs, the POH profile of FISHR was the most similar to the profile expected when the estimated endpoints of the called segments are perfect (Figure 8A); FISHR had levels of POH in the two flanking regions (“downstream” and “upstream”) very close to that between pairs of random individuals, indicating very little under-extension, and it had ~0 POH in quarters 1 through 4, indicating very little overextension. rIBD was very precise at finding segments that were truly IBD (~0 POH in quarters 1 through 4), but as predicted, it tended to under-extend the IBD segments much more than any of the other programs (low POH in the flanking regions). On the other hand, HaploScore tended to overextend true IBD segments, as indicated by its higher POH in the first and fourth quarters. GERMLINE tended to both overextend called segments and under-extend them, especially at the beginning of called segments. Supplementary Figure S23 illustrates the same POH analysis but instead uses a minimum of 1 cM for called IBD segments.

**Figure 5.**
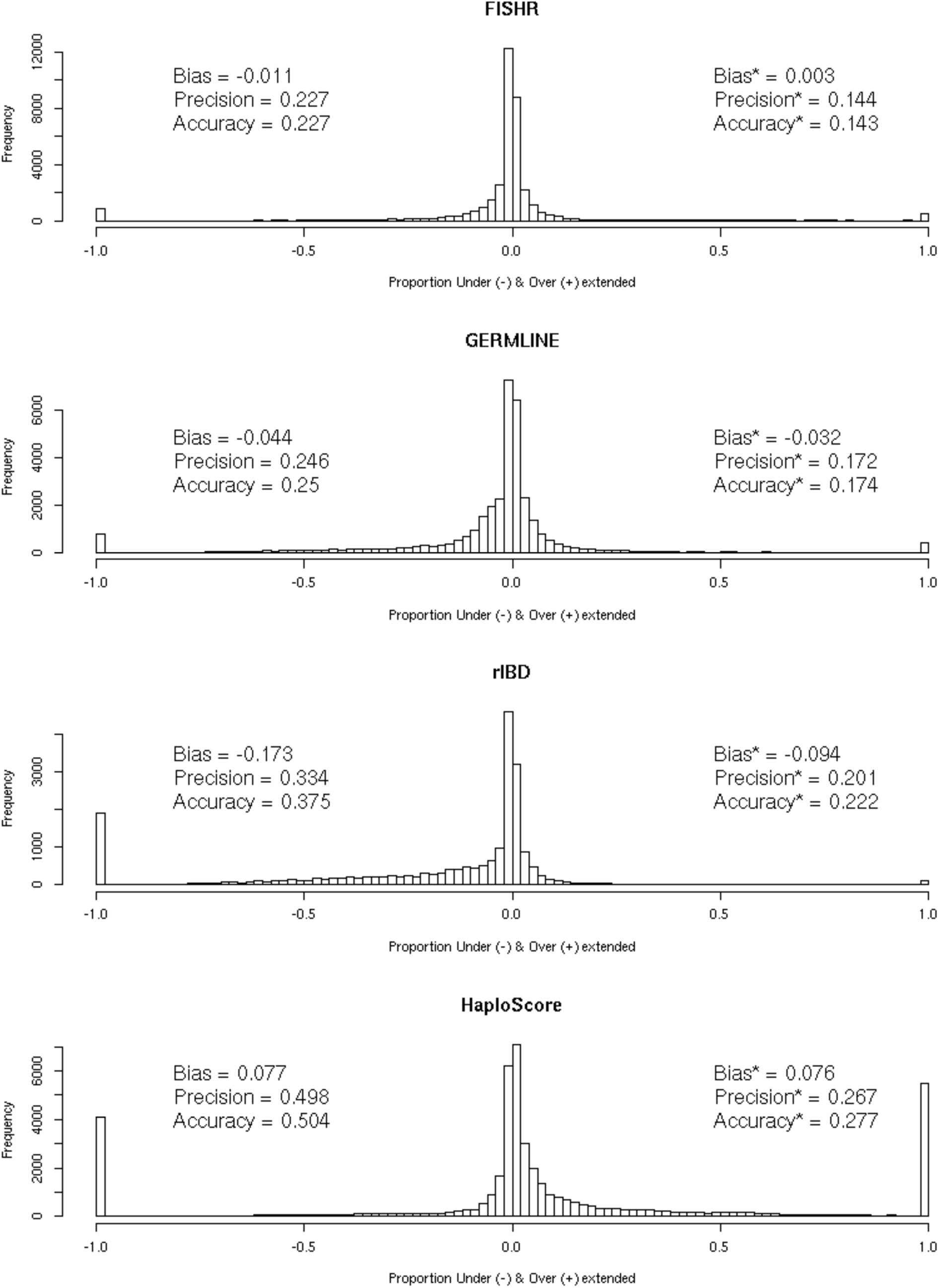
Histograms displaying the distributions of the proportional under‐ and overextension for each called IBD segment for FISHR, GERMLINE, rIBD, and HaploScore, with the bias, precision, and accuracy observed for each program. Results were found using a minimum of 3 cM for called segments and 1.5 cM for true IBD segments. All called segments with no corresponding true IBD segments (the entire segment was overextended) were classified as 1, and all true segments with no corresponding called segments (the entire “called” segment was underextended) were classified as ‐1. Results listed on the left sides on the histograms include these false positive and false negative calls while the results listed on the right sides of histograms marked with a * only included the called segments which had a corresponding true IBD segment.

**Figure 6.**
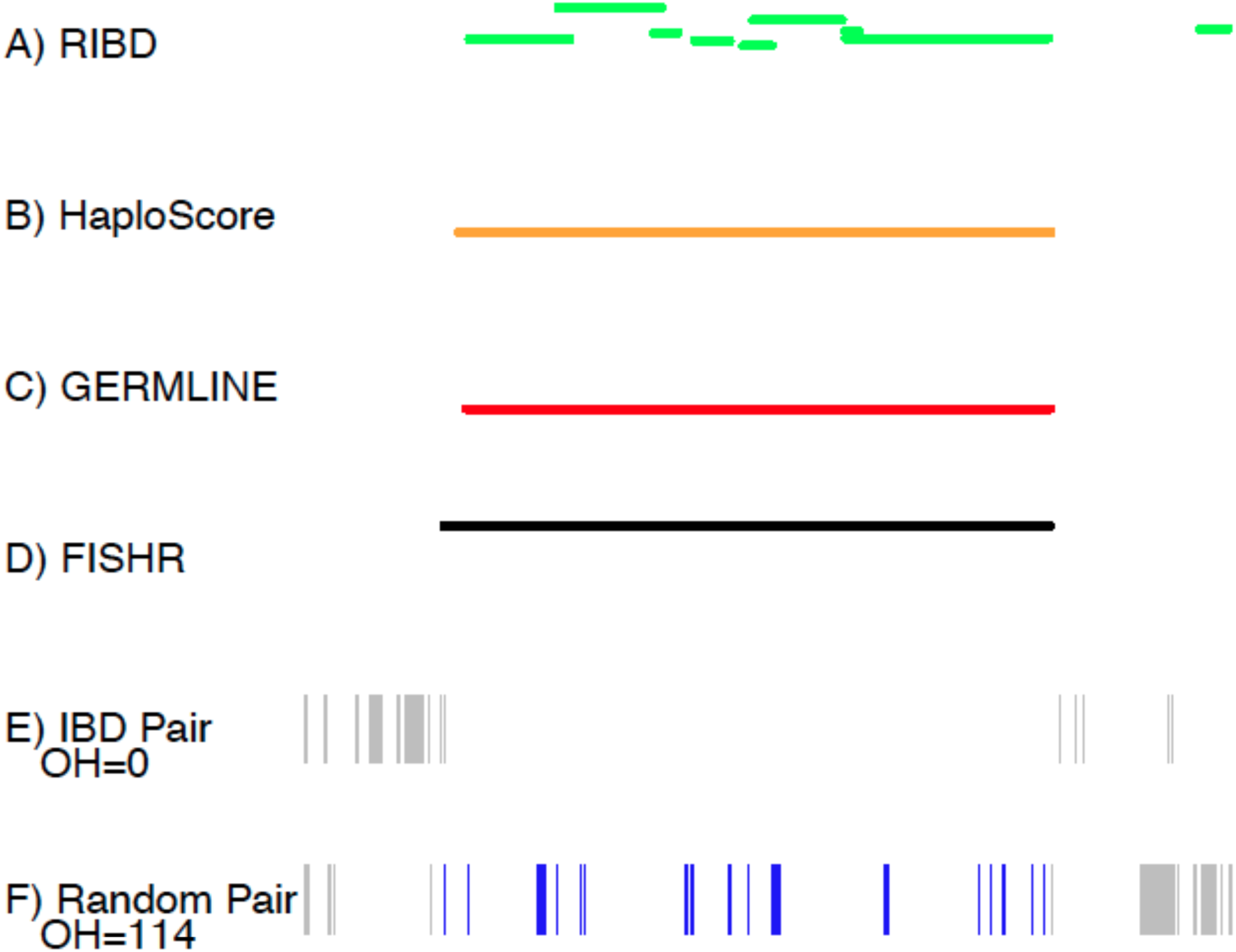
An example of called IBD segments between two individuals in the UK10K dataset, from (A) rIBD, (B) HaploScore, (C) GERMLINE, and (D) FISHR, with (E) opposite homozygous SNPs (OH) occurring for that pair of individuals in and surrounding the FISHR called IBD segment with the number of OH within the called segment listed, and (F) OH occurring in a random pair of individuals at the same location of the called IBD segment with the number of OH listed. (Note that rIBD can call two individuals as IBD 2 at some locations, i.e. sharing two IBD haplotypes; hence the overlapping segments shown for that program.)

**Figure 7.**
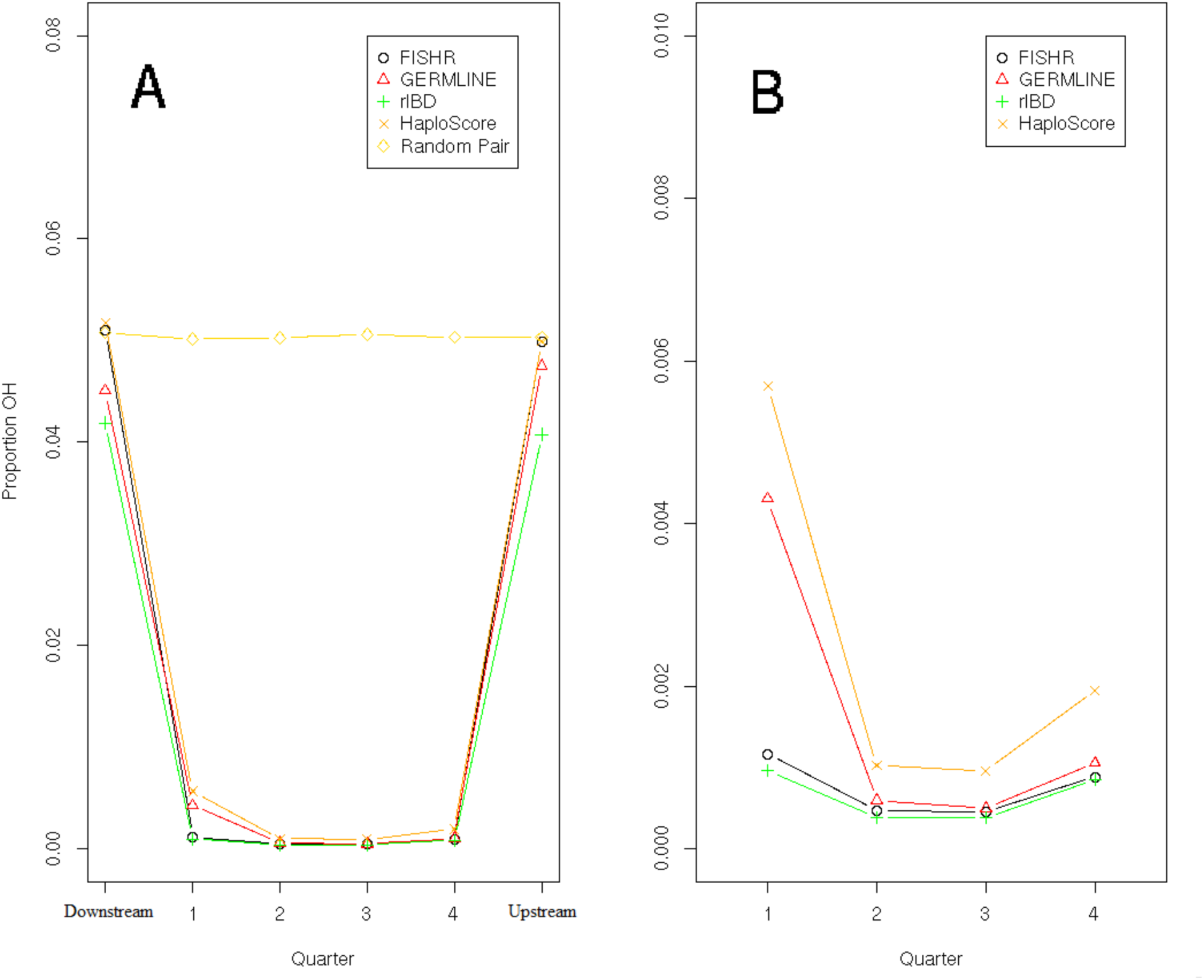
Results of the analysis of proportion of opposite homozygosity (OH) in (A) four quarters of called IBD segment and the two flanking regions and in (B) just the four quarters of the called IBD segments for FISHR (o), GERMLINE (Δ), rIBD (+), HaploScore (x), and random individuals at the same location of called IBD (◊) where called IBD segments were a minimum of 3 cM.

**Figure 8.**
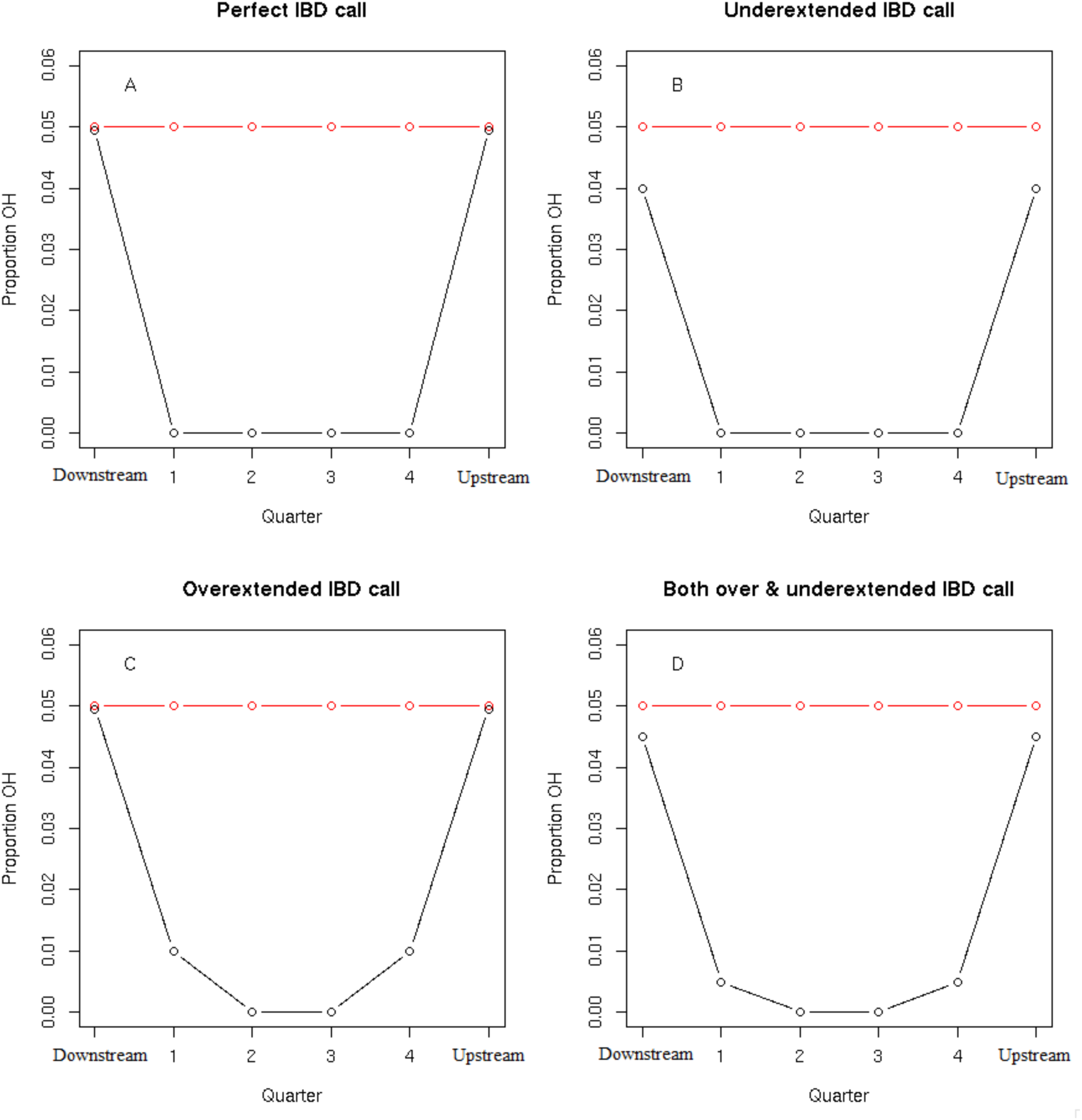
Examples that summarize the proportion of opposite homozygosity (POH) calculated from 4 quarters within and the two flanking regions around each called IBD segment, with the POH for the called segment in black and the POH of segments from random individuals at the same location in the genome presented in red. A program that makes every IBD call perfectly from perfectly genotyped data (A) would have no OH in quarters one through four and the same POH as random segments in the flanking regions. A program that underextends calls (B) would have no OH in quarters one through four and lower POH in the flanking regions than random segments. A program that overextends calls (C) would have positive POH in the first and fourth quarters and the same POH in flanking regions as random segments. A program that both under and overextends IBD calls (D) would display both increased POH in quarters one and four and decreased POH in the flanking regions.

## Discussion

We developed FISHR as an alternative method to detect segments of the genome shared IBD between pairs of individuals in a sample measured on genome-wide SNP data. Our goal was to develop a program that would be fast enough to be utilized with very large SNP datasets and be more accurate than existing programs at detecting IBD segments and their true endpoints. As we demonstrated using simulated data where true IBD status was known, FISHR performs as well or better than all competitor programs in terms of PPV and sensitivity for detecting long IBD segments, while slightly worse than rIBD but better than GERMLINE and HaploScore at detecting short IBD segments. Furthermore, as we demonstrated in both simulated and real data, FISHR is substantially more accurate than any existing program at estimating the correct endpoints of IBD segments. Accurately estimating these endpoints is important for several reasons. First, the length of IBD segments is relevant to many parameters of interest in population genetics (time to recent common ancestor, effective population size, population bottlenecks, etc.); systematic biases in estimating these lengths can lead to incorrect conclusions regarding these and other parameters. Second, phasing and imputation (Kong et al. 2008) based on IBD segments can be affected by the accuracy of the endpoints, with under‐ and overextensions of IBD segments causing regions to be incorrectly imputed or phased. Finally, in calculating genome-wide relatedness using IBD segments (Browning and Browning 2013), programs that tend to overextend IBD calls will lead to systematically inflated relatedness, and those that tend to underextend IBD calls to deflated relatedness.

Despite the computationally efficient, deterministic algorithm FISHR uses to call candidate segments (see *Methods*), FISHR remains surprisingly accurate. It is fast enough to be used on very large SNP datasets (e.g., 20,000-50,000 individuals), running two to five times slower than GERMLINE but running over a thousand times faster than rIBD and HaploScore at large sample sizes. One practical downside of FISHR is that it requires much more RAM than its competitors. This is because FISHR attempts to stitch together long called segments that are separated by a small number of SNPs, which may represent erroneously split IBD segments (although FISHR may subsequently break up some of these consolidated segments if the data suggests the full segment is not IBD). To accomplish this, FISHR must pull all the candidate segments from GERMLINE into RAM to sort them, making its memory overhead high compared to programs such as GERMLINE that simply stream data. However, given that the price of RAM is plummeting, and that the RAM capacity of many high-performance computers (e.g., 1 Tb) is already sufficiently large for FISHR to be applied on samples of ~100,000, we do not see this as a major impediment to using the program. Nevertheless, we have developed a version of FISHR (accessed using the *-low_ram* flag) that uses a negligible amount of RAM at the cost of failing to stitch together called segments that are erroneously split. The accuracy of this version of FISHR is only slightly degraded compared to the default version.

Another limitation of FISHR vis-à-vis rIBD is that, using the approach we presented here, it cannot call regions that are greater than IBD 1 – i.e., where more than one IBD segment exists at the same location between individuals. For example, ~25% of regions between siblings are expected to be IBD 2, meaning both haplotypes are IBD. FISHR (as well as GERMLINE) would call these regions as IBD 1, whereas rIBD can call these regions as IBD 2 (or greater). We have incorporated a method for detecting such multi-IBD states into FISHR (by post-processing GERMLINE segments found using the *-haploid* flag), but because such IBD 2+ situations are extremely rare among unrelated individuals (occurring at a rate proportional to the square of relatedness, or ~0.0001 for IBD 2 vs. 0.01 for IBD 1 in typical datasets of nominally unrelated individuals), the benefit of these additional called segments did not outweigh the cost in missing truly IBD segments incurred by post-processing data called using *-haploid* in GERMLINE. Nevertheless, the standard version’s limitation to detecting IBD 1 must be kept in mind when working with highly related samples.

## Conclusion

With increasingly large whole-genome SNP datasets being accumulated, it is important to have a method for detecting IBD segments that is both accurate and efficient. We introduced a program, FISHR, that accomplishes both, and that is particularly accurate at determination of the correct endpoints of IBD segments. We demonstrated these properties using simulations, and confirmed these conclusions using a novel approach on real sequence data from the UK10K project. Due to the number of pairwise comparisons that must be made in IBD detection, computationally intensive programs such as rIBD and HaploScore cannot be easily run on datasets of more than ~10,000 individuals. FISHR is a more accurate alternative to GERMLINE as an IBD detection program on large datasets, with only a modest increase in runtime.

## Methods

### Description of the FISHR algorithm

FISHR is written in C++ and is available freely for download at

http://matthewckeller.com/html/program_code.html. FISHR utilizes GERMLINE (described in detail by Gusev et al. 2009), as an initial screen to quickly detect candidate segments. In particular, in the results presented here, we used the *-hextend* method in GERMLINE, which incorporates information on phased mismatches and which we found to be the most accurate of the three alternative methods (*-h extend, ‐w extend*, and *-haploid*) GERMLINE uses. FISHR then further refines the called segments as follows. First, because two long IBD calls that are separated by a short distance may actually be a single contiguous IBD segment that was artificially broken apart in GERMLINE due to phase or SNP call errors, FISHR stitches together segments separated by a user-defined number of SNPs (-gap). Next, FISHR finds the locations of IEs for all called segments. To do this, FISHR finds the longest exact match between either of the two phased haplotypes of the first person and either of the two phased haplotypes of the second person (a total of four possible combinations), starting at the first SNP of the called segment. An IE occurs at the first mismatching SNP after the exact match ends. FISHR then finds the next longest exact match between any of the four possible combinations of phased haplotypes, starting from the SNP following the previous IE, and extends until the next mismatching SNP is encountered. This process is continued until the end of the called segment.

IEs represent locations along a candidate segment that are potentially inconsistent with IBD inheritance. Some IEs are expected by chance due to SNP and phase errors even in truly IBD segments. However, too many IEs within a particular region are a likely signal that the segment is not IBD in that area and that the segment should be truncated (if near an endpoint of the segment) or split into two (if in the middle of the segment). To determine such called segment endpoints, FISHR calculates a moving average (MA) of IEs centered at each SNP within a user-defined window (using the *-window* flag) of SNPs, as outlined in Figure 9. FISHR then starts at the center of the called IBD segment and moves towards each endpoint until it reaches the first SNP with a MA value greater than the user-defined maximum (*-empmathreshold*), as shown in Figure 10. These points signal the endpoints of a called segment. Note that in addition to trimming the segment ends, this process can split a GERMLINE candidate segment into two or more shorter segments. Moreover, if the flag *-count gap errors* is set to TRUE, as it is by default, segments that had been stitched together from the first step can broken up again at this stage if enough IEs are clustered near the gap. Because segments that are too short, in terms of either number of SNPs or cM distance, are increasingly likely to be false positives, FISHR then drops segments shorter than user-defined thresholds of both SNP and cM length (using the *-min snp* and *-mincm* flags, respectively). The final process FISHR performs is to calculate the total proportion of SNPs that are IEs (PIE) within each segment. Too many IEs scattered across the entire length of a segment are a signal that the whole segment is unlikely to be IBD. Thus, if the PIE of a segment is greater than the value supplied in the *-emp_pie_threshold* argument, the segment is dropped.

**Figure 9.**
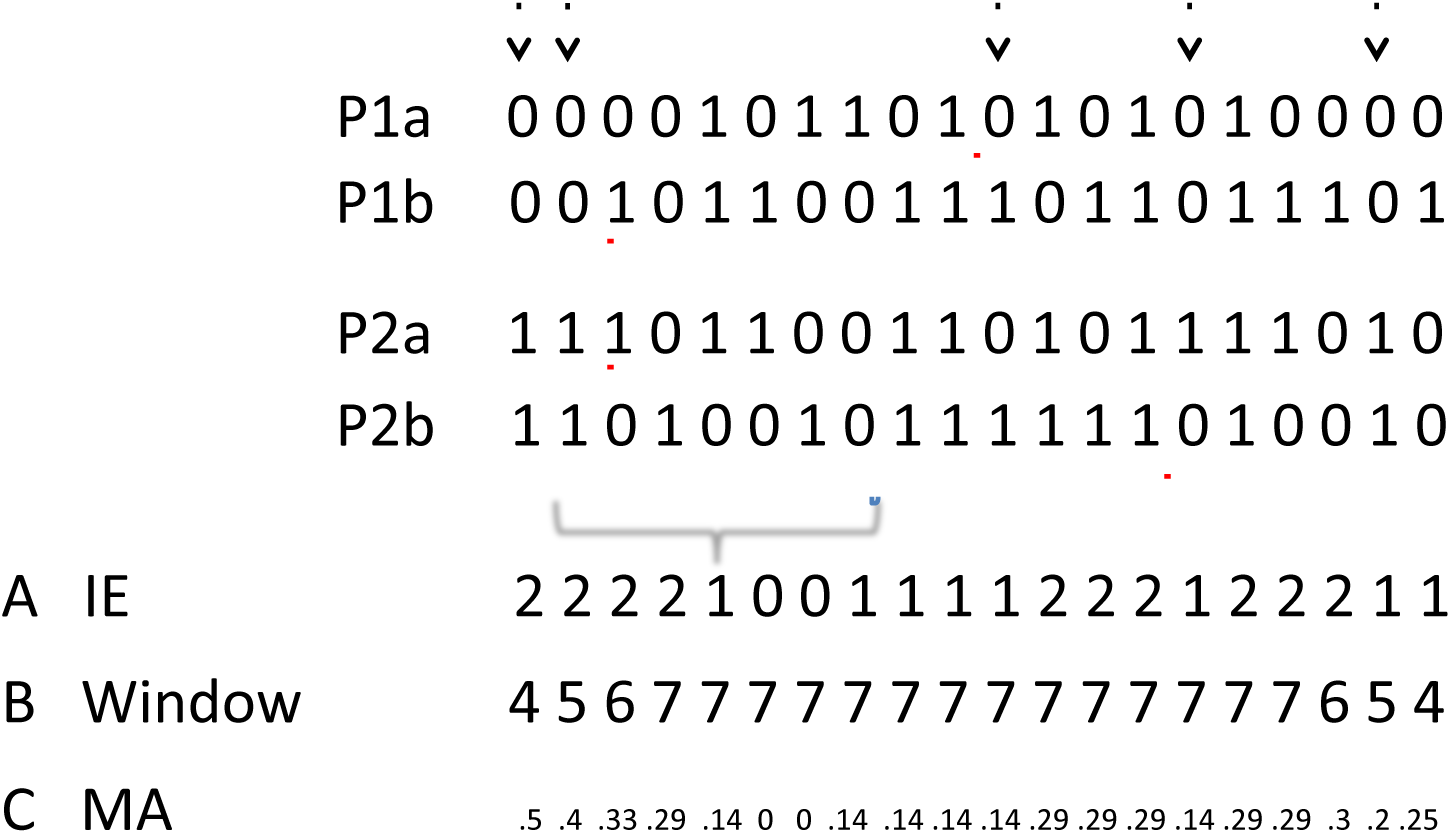
Calculating the moving average (MA) of implied errors (IE) of a potential IBD call between two individuals, P1 & P2. The red underlined segments indicate the called haplotypes, and the arrows designate where IEs occur in the call. Using a moving window size of 7, line A displays the number of IEs within the window for each given SNP, line B displays the window size (which is truncated at each end of the “chromosome”), and line C displays the MA for each SNP.

**Figure 10.**
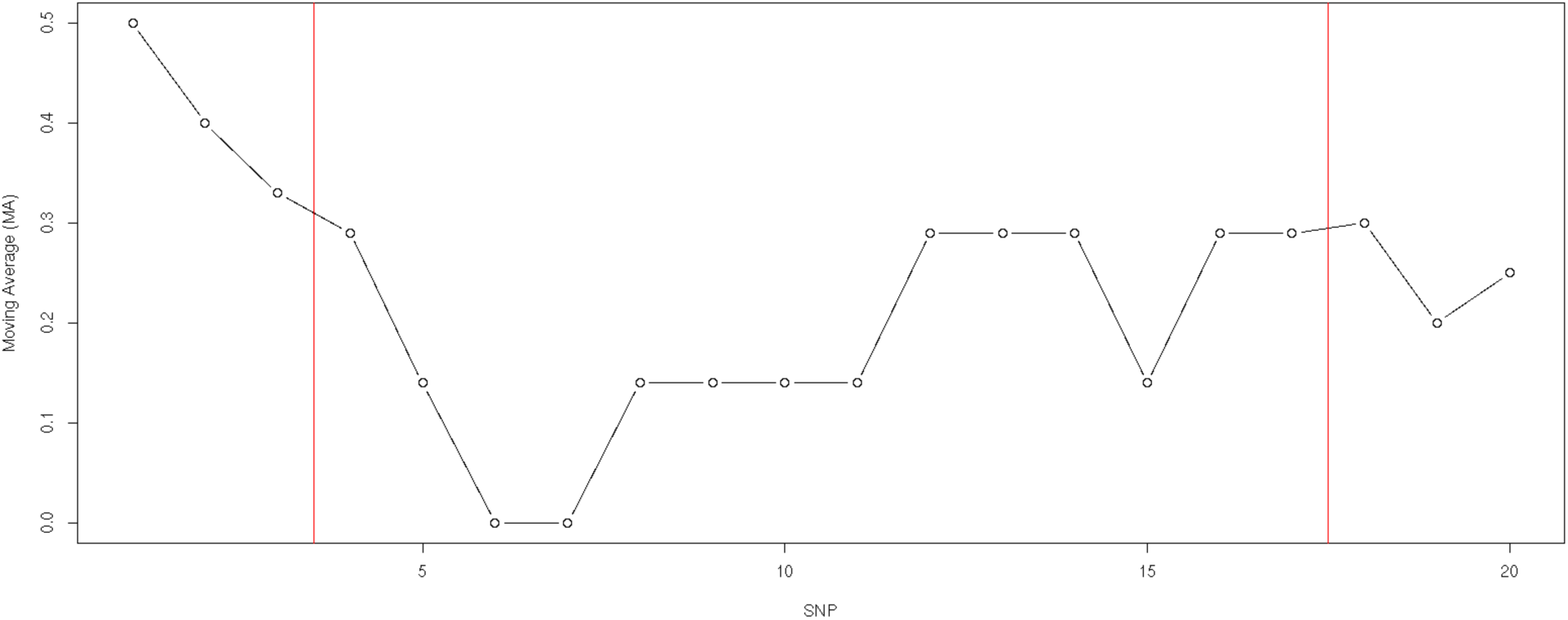
An example how FISHR utilizes the moving average of implied errors (MA) calculated for each SNP to determine endpoints of a called IBD segment. In this case the maximum allowed MA is 0.3. The red vertical line denotes the location of where the MA increases above the 0.3 threshold, leaving the called segment to consist of the SNPs between the red lines.

Because values of PIE and MA depend on the quality of SNP calls and phasing in the data at hand, the thresholds for these values require careful consideration by users. The approach we recommend and that we used here was to identify long stretches (>8 cM) of the genome where no opposite homozygotes occurred between pairs of individuals (this can be accomplished using GERMLINE *-w_extend* flag without the *-h_extend* flag). Because information on phase was not used in calling these segments, they are not biased to be in regions where phasing is more difficult. We then found the distribution of the PIE and maximum MA values calculated from the middlemost 50% of these segments, which can be assumed with high confidence to be truly IBD. We compared these distributions to distributions of PIE and maximum MA values calculated from segments matched in location to the likely IBD segments but that were between random pairs of individuals. The PPV and sensitivity that will result from any choices of PIE and MA thresholds can be estimated by how well those thresholds separate these distributions, and thus thresholds can be chosen that lead to a desired PPV-sensitivity combination. We have supplied a utility (*gl_parameter_finder*) for accomplishing this step along with the FISHR download.

### Simulated Sequence and SNP Data

We simulated genotypic data using the sequence simulator HAPGEN2 (Su et al. 2011), which simulated haplotypes by conditioning on a reference set of population haplotypes (here, the 1000 Genomes Project (Clarke et al. 2012) European ancestry (CEU) haplotypes of chromosome 15) and created a new population by combining haplotypes according to a fine-scaled recombination rate map (from deCODE; Kong et al. 2010). Here, we defined the effective population sizes as 11,418 and the sample size (defined as “controls” in HAPGEN2) as 28,000. For computational efficiency, we created 13 independent datasets of 1,000 individuals each and averaged all results across these 13 replicates. The data had LD, haplotype diversity, and allele frequency distributions that mimic those in the initial set of haplotypes.

We used the perfectly phased, simulated sequence data with no errors obtained from HAPGEN2 to obtain “true IBD segments.” Because no program exists to our knowledge that tracks IBD status between pairs of haplotypes, we defined true IBD segments as perfectly matching haplotypes that spanned a desired cM threshold (0.25, 0.5, 1.5, or 3, depending on the analysis).

To increase computational efficiency and to ensure that rare mutations that arose on a haplotype since the common ancestor did not cause a true IBD segment to be missed, we pruned this sequence data to have MAF > .05, resulting in a density of ~1 variant per 1000 base pairs. To create data that mimicked post-quality-control SNP data on existing platforms, we then extracted SNPs pseudo-randomly such that the MAF distribution was about uniform and the density of SNPs was one per 6,750 base pairs (corresponding to ~400,000 SNPs genomewide). To simulate SNP call errors, we randomly changed one allele to its alternative allele at a rate of 0.2%, in the middle of what has been found empirically for SNP calls (Steemers and Gunderson 2007; Teo et al. 2007; Korn et al. 2008; Hong et al. 2012). Finally, we unphased the SNP data and rephrased it using SHAPEIT2 (Delaneau et al. 2012).

### Real Sequence Data

We also compared performance of the IBD detection algorithms using the UK10K ALSPAC sequence data on 1,872 unrelated individuals (The UK10K Consortium 2015). In this data, we utilized 4 subchromosomes (5q, 9q, 14q, and 20q) and removed markers with less than a 1% MAF, markers in violation of Hardy-Weinberg equilibrium with p-values of less than 0.0001, and markers that contained missing data for any individuals. We then extracted SNPs that were on the Illumina 650K SNP panel (21,802 markers for subchromosome 5q, 13,716 markers for subchromosome 9q, 16,199 markers for subchromosome 14q, and 6,307 markers for subchromosome 20q) and phased this data using SHAPEIT2 for calling segments using each program. We retained the remaining markers not in the SNP data (an average of one marker per 3,000 base pairs) as a holdout sample to calculate the proportion of opposite-homozygote SNPs within called segments.

### Running the four IBD detection programs

We ran FISHR, GERMLINE, rIBD, and HaploScore on the simulated SNP data that was phased using SHAPEIT2, varying input parameters to determine the optimal parameters for discovering IBD segments with minimum lengths both of 1 and 3 cM for each program (optimal parameters for finding IBD with a minimum of 3 cM bolded). For FISHR, we varied the candidate segment detection parameters (using GERMLINE) ***–h_extend*** vs. *–w_extend, ‐bits* (30, 45, **60**, 75, or 90), *-err_het* (0, **1**, 2, 3), *-errhom* (0, **1**, 2, 3), as well as the FISHR-specific parameters *-gap* (0, 1, or **30**), *-count_gap_errors* (**TRUE** or FALSE), *-emp_ma_threshold* (0.025, **0.045**, 0.065, 0.085), and *-emp_pie_threshold* (0.005, **0.015**, 0.025). For GERMLINE, we compared both the ***-h_extend*** vs. *–w extend* options and varied *–bits* (30, 45, **60**, 90, 120, 150), *-err het* (0, **1**, 2, 3), and *-err hom* (0, **1**, 2, 3). For rIBD, we varied *–ibdlod*(**1**, 2, 3, 4, 5, 6), *-overlap* (100, **157**, 200), *-window* (7,500, **10,000**, 12,500), *-scale* (2.5, 3, **3.16**, 3.5), and *-trim* (11, **16**, 21). Finally, for HaploScore, we varied the candidate segment detection parameters (using GERMLINE) of ***-h_extend*** vs. *-w_extend, ‐bits* (**30**, 45, 60, 90, 120), *-err_het* (0, **1**, 2, 3), and *-err_hom* (0, **1**, 2, 3), and then varied the *-switch_error* (**0.0005**, 0.001, 0.0015, 0.01), *–snp_error* (**0.0006**, 0.00125, 0.0025, 0.01), and HaploScore thresholds (**1**, 3, 5, 7, 9, 11, 13, 15) in HaploScore. For each program, we plotted the PPV and sensitivity, as shown in Figure 4, and the combination closest to perfect performance (Sensitivity=1 and PPV=1) was kept as the optimal for that specific program. The exact command lines used for each program with these optimal parameters are included in Supplemental Table S2. For Figures 3 and 4, we kept constant all the optimal parameters for each program other than the parameter that most influenced the PPV-sensitivity tradeoff. In particular, we varied *–emp_ma_threshold* for FISHR, *–ibdlod* for rIBD, and the *–bits* argument for GERMLINE and HaploScore.

## Data Access

MGS dataset(s) used for the analyses described in this manuscript were obtained from dbGaP found at http://www.ncbi.nlm.nih.gov/gap through dbGaP study accession numbers phs000167. This dataset was provided by Alan R. Sanders, M.D. CARDIA dataset(s) used for the analyses described in this manuscript were obtained from dbGaP found at

http://www.ncbi.nlm.nih.gov/gap through dbGaP study accession numbers phs000309. The ARIC datasets used for the analyses described in this manuscript were obtained from dbGaP found at http://www.ncbi.nlm.nih.gov/gap through dbGaP study accession numbers phs000090. The GENEVA datasets used for the analyses described in this manuscript were obtained from dbGaP found at http://www.ncbi.nlm.nih.gov/gap through dbGaP study accession numbers phs000091. Simulated data, scripts to evaluate IBD detection, and FISHR can be downloaded from our personal website, http://matthewckeller.com/html/.

## Acknowledgements

The authors thank Nathan Lapinski, Teresa deCandia, and Rasool Tahmasbi for their help in coding, ideas, and writing.

The Atherosclerosis Risk in Communities Study was carried out as a collaborative study supported by National Heart, Lung, and Blood Institute contracts (HHSN268201100005C, HHSN268201100006C, HHSN268201100007C, HHSN268201100008C, HHSN268201100009C, HHSN268201100010C, HHSN268201100011C, and HHSN268201100012C). The authors thank the staff and participants of the ARIC study for their important contributions. Funding for GENEVA was provided by National Human Genome Research Institute grant U01HG004402 (E. Boerwinkle).

Funding support for the GWAS of Gene and Environment Initiatives in Type 2 Diabetes was provided through the NIH Genes, Environment and Health Initiative [GEI] (U01HG004399).

The human subjects participating in the GWAS derive from The Nurses’ Health Study and Health Professionals’ Follow-up Study and these studies are supported by National Institutes of Health grants CA87969, CA55075, and DK58845. Assistance with phenotype harmonization and genotype cleaning, as well as with general study coordination, was provided by the Gene Environment Association Studies, GENEVA Coordinating Center (U01 HG004446). Assistance with data cleaning was provided by the National Center for Biotechnology Information. Funding support for genotyping, which was performed at the Broad Institute of MIT and Harvard, was provided by the NIH GEI (U01HG004424).

## Author contributions

M.C.K. and M.J. conceived the concept for the program. U.L. and P.S.P. coded the program. D.W.B and M.C.K. analyzed and compared results from the programs and wrote the manuscript.

## References

Browning BL, Browning SR. 2011. A fast, powerful method for detecting identity by descent. The American Journal of Human Genetics 88: 173–182.

Browning SR, Browning BL. 2012. Identity by descent between distant relatives: detection and applications. Annual review of genetics 46: 617–633.

Browning SR, Browning BL. 2013. Identity-by-descent-based heritability analysis in the Northern Finland Birth Cohort. Human genetics 132: 129–138.

Clarke L, Zheng-Bradley X, Smith R, Kulesha E, Xiao C, Toneva I, Vaughan B, Preuss D, Leinonen R, Shumway M. 2012. The 1000 Genomes Project: data management and community access. Nature methods 9: 459–462.

Delaneau O, Marchini J, Zagury J-F. 2012. A linear complexity phasing method for thousands of genomes. Nature methods 9: 179–181.

Durand EY, Eriksson N, McLean CY. 2014. Reducing pervasive false-positive identical-by-descent segments detected by large-scale pedigree analysis. Molecular biology and evolution: msu151.

Gusev A, Lowe JK, Stoffel M, Daly MJ, Altshuler D, Breslow JL, Friedman JM, Pe'er I. 2009. Whole population, genome-wide mapping of hidden relatedness. Genome research 19: 318–326.

Haldane J. 1919. The combination of linkage values and the calculation of distances between the loci of linked factors. J Genet 8: 299–309.

Hong H, Xu L, Liu J, Jones WD, Su Z, Ning B, Perkins R, Ge W, Miclaus K, Zhang L. 2012. Technical reproducibility of genotyping SNP arrays used in genome-wide association studies. PLoS One 7: e44483.

Keller MC, Visscher PM, Goddard ME. 2011. Quantification of inbreeding due to distant ancestors and its detection using dense single nucleotide polymorphism data. Genetics 189: 237–249.

Kong A, Masson G, Frigge ML, Gylfason A, Zusmanovich P, Thorleifsson G, Olason PI, Ingason A, Steinberg S, Rafnar T. 2008. Detection of sharing by descent, long-range phasing and haplotype imputation. Nature genetics 40: 1068–1075.

Kong A, Thorleifsson G, Gudbjartsson DF, Masson G, Sigurdsson A, Jonasdottir A, Walters GB, Jonasdottir A, Gylfason A, Kristinsson KT. 2010. Fine-scale recombination rate differences between sexes, populations and individuals. Nature 467: 1099–1103.

Korn JM, Kuruvilla FG, McCarroll SA, Wysoker A, Nemesh J, Cawley S, Hubbell E, Veitch J, Collins PJ, Darvishi K. 2008. Integrated genotype calling and association analysis of SNPs, common copy number polymorphisms and rare CNVs. Nature genetics 40: 1253–1260.

Palamara PF, Lencz T, Darvasi A, Pe’er I. 2012. Length distributions of identity by descent reveal fine-scale demographic history. The American Journal of Human Genetics 91: 809–822.

Powell JE, Visscher PM, Goddard ME. 2010. Reconciling the analysis of IBD and IBS in complex trait studies. Nat Rev Genet 11: 800–805.

Schizophrenia Working Group of Psychiatric Genomics Consortium, Ripke S, Neal BM et al. 2014. Biological insights from 108 schizophrenia-associated genetic loci. Nature 511: 421–427.

Setty MN, Gusev A, Pe'er I. 2011. HLA type inference via haplotypes identical by descent. Journal of Computational Biology 18: 483–493.

Soi S, Scheinfeldt L, Lambert C, Hirbo J, Ranciaro A, Thompson S, Bodo J, Froment A, Ibrahim M, Juma A. 2011. Demographic histories of African hunting-gathering populations inferred from genome-wide SNP variation. In International Congress of Human Genetics/American Society of Human Genetics meeting Montreal, Canada.

Steemers FJ, Gunderson KL. 2007. Whole genome genotyping technologies on the BeadArray™ platform. Biotechnology journal 2: 41–49.

Su Z, Marchini J, Donnelly P. 2011. HAPGEN2: simulation of multiple disease SNPs. Bioinformatics 27: 2304–2305.

Sudlow C, Gallacher J, Allen N, Beral V, Burton P, Danesh J, Downey P, Elliott P, Green J, Landray M. 2015. UK Biobank: an Open Access resource for identifying the causes of a wide range of complex diseases of middle and old age. PLoS medicine 12: 1–10.

Teo YY, Inouye M, Small KS, Gwilliam R, Deloukas P, Kwiatkowski DP, Clark TG. 2007. A genotype calling algorithm for the Illumina BeadArray platform. Bioinformatics 23: 2741–2746.

The UK10K Consortium 2015. The UK10K project identifies rare variants in health and disease. Nature 526: 82–90.

Vacic V, Ozelius LJ, Clark LN, Bar-Shira A, Gana-Weisz M, Gurevich T, Gusev A, Kedmi M, Kenny EE, Liu X. 2014. Genome-wide mapping of IBD segments in an Ashkenazi PD cohort identifies associated haplotypes. Human molecular genetics 23: 4693–4702.

